# Rodent mnemonic similarity task performance requires the prefrontal cortex

**DOI:** 10.1101/2020.08.05.238725

**Authors:** Sarah A. Johnson, Sabrina Zequeira, Sean M. Turner, Andrew P. Maurer, Jennifer L. Bizon, Sara N. Burke

## Abstract

Mnemonic similarity task performance, in which a known target stimulus must be distinguished from similar lures, is supported by the hippocampus and perirhinal cortex and is known to decline in advanced age. Interestingly, disrupting hippocampal activity leads to mnemonic discrimination impairments when lures are novel, but not when they are familiar. This observation suggests other brain structures can support discrimination abilities as stimuli are learned. The prefrontal cortex (PFC) is critical for retrieval of remote events and executive functions, such as working memory, and is also particularly vulnerable to dysfunction in aging. Importantly, the medial PFC is reciprocally connected to the perirhinal cortex and neuron firing in this region coordinates communication between lateral entorhinal and perirhinal cortices to presumably modulate hippocampal activity. This anatomical organization and function of the medial PFC suggests that it contributes to mnemonic discrimination; however, this notion has not been empirically tested. In the current study, young adult male and female F344 x Brown Norway F1 hybrid rats were trained on a rodent object-based mnemonic similarity task, and surgically implanted with guide cannulae targeting prelimbic and infralimbic regions of the medial PFC. Prior to mnemonic discrimination tests, rats received PFC infusions of the GABA_A_ agonist muscimol. Analyses of expression of the neuronal activity-dependent immediate-early gene *Arc* in medial PFC and adjacent cortical regions confirmed muscimol infusions led to neuronal inactivation in the infralimbic and prelimbic cortices. Moreover, muscimol infusions in PFC impaired mnemonic discrimination performance relative to the vehicle control across all testing blocks when lures shared 50-90% feature overlap with the target. Thus, in contrast to prior results from rats given hippocampal muscimol infusions, PFC inactivation impaired target-lure discrimination regardless of the novelty or familiarity of the lures. These findings indicate the PFC plays a critical role in mnemonic similarity task performance, but the time course of PFC involvement is dissociable from that of the hippocampus.

## Introduction

The ability to discriminate between stimuli that share features declines in advanced age. Specifically, older adults are impaired at distinguishing similar lures from previously seen target stimuli relative to young adults, as assessed by the ‘mnemonic similarity task’ (Toner et al., 2009; Holden et al., 2013; Stark et al., 2013, 2015, 2019; Reagh et al., 2016; Huffman and Stark, 2017; Trelle et al., 2017, 2020; Camfield et al., 2018). Mnemonic similarity task deficits in advanced age have been reported for older adults who are able to perform on par with young adults in standardized neuropsychological test batteries (Stark et al., 2013; Reagh et al., 2014, 2016). These data suggest that tasks assessing stimulus discrimination abilities are particularly sensitive in detecting cognitive decline and could provide a behavioral read-out of underlying neural dysfunction that accompanies aging. In support of this idea, human neuroimaging studies have shown that age-related mnemonic discrimination impairments are associated with reduced integrity of white matter afferents to the hippocampus (Yassa et al., 2010; Bennett et al., 2015; Bennett and Stark, 2016) and altered task-related activation in the perirhinal cortex (Ryan et al., 2012; Berron et al., 2018), lateral entorhinal cortex (Bakker et al., 2015; Reagh et al., 2018; Berron et al., 2019), and hippocampal dentate gyrus (DG)/CA3 (Yassa et al., 2011a, 2011b; Bakker et al., 2012, 2015; Doxey and Kirwan, 2015; Reagh et al., 2018). These data echo prior findings that cortical inputs to the hippocampus are particularly vulnerable in aging (Barnes and McNaughton, 1980; Hyman et al., 1984; Geinisman et al., 1992).

Behavioral mnemonic discrimination impairments in late adulthood are also observed in animal models. Aged rats and monkeys, like humans, are prone to false recognition and are more likely to respond as if novel objects are familiar (Burke et al., 2010). This false recognition is particularly evident when the objects share features with previously experienced stimuli (Burke et al., 2011). Further, in an object discrimination task designed to directly parallel human mnemonic similarity tasks (Stark et al., 2019), aged rats were selectively impaired in distinguishing similar objects relative to young adult rats (Johnson et al., 2017). Importantly, discrimination of objects without common features is comparable between young and old animals (Burke et al., 2011; Hernandez et al., 2015; Johnson et al., 2017).

Given CA3/DG activity has been consistently linked to mnemonic discrimination performance in both young and older adults (Kirwan and Stark, 2007; Bakker et al., 2008; Lacy et al., 2011; Yassa et al., 2011a; Motley and Kirwan, 2012; Reagh and Yassa, 2014; Doxey and Kirwan, 2015; Reagh et al., 2018), we recently tested the requirement of intact neural activity in these hippocampal regions for distinguishing similar objects in rats (Johnson et al., 2018). While disrupting activity within CA3/DG impaired discrimination performance in young adults, this effect was contingent on prior task experience. Specifically, infusions of the GABA_A_ agonist muscimol in CA3/DG impaired discrimination in a first block of tests when lure objects were novel, but not on subsequent second and third blocks of tests as lures became familiar (Johnson et al., 2018). In contrast, interference with perforant path input to the hippocampus impaired mnemonic discrimination performance across multiple tests (Burke et al., 2018b). These findings raised two points of interest. First, in agreement with data from humans, intact cortical input to the hippocampus is important for mnemonic discrimination performance. Second, as experience with procedural or contextual aspects of a task accrues, extra-hippocampal regions appear to compensate for altered activity within the hippocampal circuit.

It is well established that prefrontal cortical activity and connectivity with hippocampal and parahippocampal circuits supports memory-guided behavior across the lifespan (Gaffan, 2002; Maillet and Rajah, 2013; Eichenbaum, 2017; Barry and Maguire, 2019; Takehara-Nishiuchi, 2020). It is therefore possible that when hippocampal activity is disrupted, prefrontal-perirhinal or -entorhinal circuits guide mnemonic discrimination behavior. In support of this interpretation, prefrontal cortical neuron spiking increases coordinated activity between perirhinal and lateral entorhinal cortices (Paz et al., 2007), which could facilitate cortical-hippocampal interactions. Conversely, blocking prefrontal neural activity or prefrontal-perirhinal communication impairs the ability to retrieve previously learned hippocampal-dependent object-place associations (Hernandez et al., 2017) and odor sequence memories (Jayachandran et al., 2019). Activation of prefrontal neuronal ensembles that receive synaptic input from the perirhinal cortex is attenuated in aged rats when tested in a hippocampal-dependent object-place association task (Hernandez et al., 2018a). Further, rats with hippocampal lesions exhibit compensatory prefrontal activity that is associated with intact expression of a learned, hippocampal-dependent, contextual fear association (Zelikowsky et al., 2013). These data point to an important role for prefrontal cortices in modulating neural activity during mnemonic discrimination, yet, to date, few studies have examined frontal cortical contributions to this cognitive function (Pidgeon and Morcom, 2016; Wais et al., 2018).

The current study investigated how prefrontal cortical activity contributes to the discrimination of similar objects in a rodent mnemonic similarity task (Johnson et al., 2017, 2018). Rats were trained on task procedures, then implanted with guide cannulae targeting prelimbic and infralimbic regions of the medial prefrontal cortex. These regions were selected because of their indirect anatomical connectivity with the hippocampus through perirhinal and entorhinal cortices (Sesack et al., 1989; Jay and Witter, 1991; Condé et al., 1995; Burwell and Amaral, 1998; Vertes, 2004; Hoover and Vertes, 2007; Agster and Burwell, 2009; Eichenbaum, 2017; Hwang et al., 2018). To test the prediction that prefrontal cortical activity is necessary for task performance, rats received infusions of the GABA_A_ agonist muscimol prior to mnemonic discrimination testing. We have previously shown that muscimol infusions at these coordinates and dose block expression of the neuronal activity-dependent immediate early gene *Arc* (Hernandez et al., 2017). This finding was replicated in the current study, confirming efficacy of prefrontal cortical inactivation through GABA_A_ agonism. Prefrontal muscimol infusions impaired mnemonic discrimination of a target object from both distinct and similar lure objects, with the effect most pronounced for similar lures. Further, in contrast to effects of hippocampal muscimol infusions (Johnson et al., 2018), disrupting prefrontal activity impaired performance irrespective of prior task experience.

## Methods

### Subjects

Seven young adult Fischer 344 x Brown Norway F1 hybrid rats (NIA, Taconic; 4-6 m at arrival) were used as subjects and completed the study in two cohorts (cohort 1: 3 males, cohort 2: 2 males, 2 females). Rats were single-housed in standard Plexiglas cages and maintained on a 12-h reversed light/dark cycle with lights off at 8:00 am. Experimental manipulations took place during the dark portion of the cycle, 5-7 days per week at the same time each day. Rats were given 7 days to acclimate to the housing facility upon arrival and were handled for a minimum of 4 days prior to beginning experiments. Rats were placed on a restricted feeding protocol after this period of acclimation, receiving 20±5 g moist chow (∼39 kcal; Teklad LM-485, Harlan Labs) daily after completing behavioral training. Water was provided *ad libitum*. Behavioral shaping began once rats reached 85% of their initial baseline body weights, which provided appetitive motivation to perform the tasks. All procedures were in accordance with the NIH *Guide for the Care and Use of Laboratory Animals* and approved by the Institutional Animal Care and Use Committee at the University of Florida.

### Habituation and procedural training

Procedures for habituation and procedural training were similar to those previously described (Johnson et al., 2017, 2018). Rats were habituated to eating small pieces of Froot Loop cereal (Kellogg’s; Battle Creek, MI) or marshmallows (Medley Hills Farm, Medina, OH), which served as food rewards throughout experiments, by providing pieces with daily chow for 1-3 days. Rats were then habituated to the task apparatus by encouraging foraging for scattered food reward pieces for 1-3 days. Tasks were carried out in a U-shaped maze built on a low table (surface 148 cm x 76 cm) out of plastic building bricks (DUPLO®, LEGO®, Enfield, CT; **Fig.1A**), similar to mazes previously used in our laboratory (Hernandez et al., 2015, 2017; Johnson et al., 2017; Maurer et al., 2017; Burke et al., 2018b; Hernandez et al., 2018a; Johnson et al., 2018; Kreher et al., 2019). Rats were trained in daily sessions to alternate between a designated start area in one arm and choice platform in the opposite arm (**Fig.1A**) by providing food reward in each location. After reaching a criterion of 32 alternations within a single daily session of 20 min or less, rats were trained on forced-choice continuous object discrimination task procedures with a pair of ‘standard’ unrelated objects (0% feature overlap; **Fig.1B**). Rats received additional training with a pair of perceptually distinct LEGO objects (38% feature overlap; **Fig.1B**). Detailed description of feature overlap calculations is provided in a previous publication from our lab (Johnson et al., 2017). For each object pair, one object served as the target (S+), placed over the baited food well, while the alternate object was a lure (S-), placed over the empty food well. Training proceeded in daily sessions of 32 trials. In each trial, rats exited the start area, traversed the maze to the choice platform, and displaced one of the two objects covering the food wells. If the target (S+) was selected, rats consumed the reward and returned to the start area to receive a second reward. If the lure (S-) was selected, both objects and the food reward were quickly removed from the choice platform and the rat did not receive a reward in the start area. The side of the baited food well on the choice platform was pseudo-randomized across trials to provide an equal number of left- and right-well trials over the course of each session. For each training object pair, the object serving as the target was counterbalanced across rats. Training was considered complete when rats reached a criterion of ≥26 correct responses out of 32 trials (≥81.3%) within a session for 2 consecutive days.

**Figure 1.**
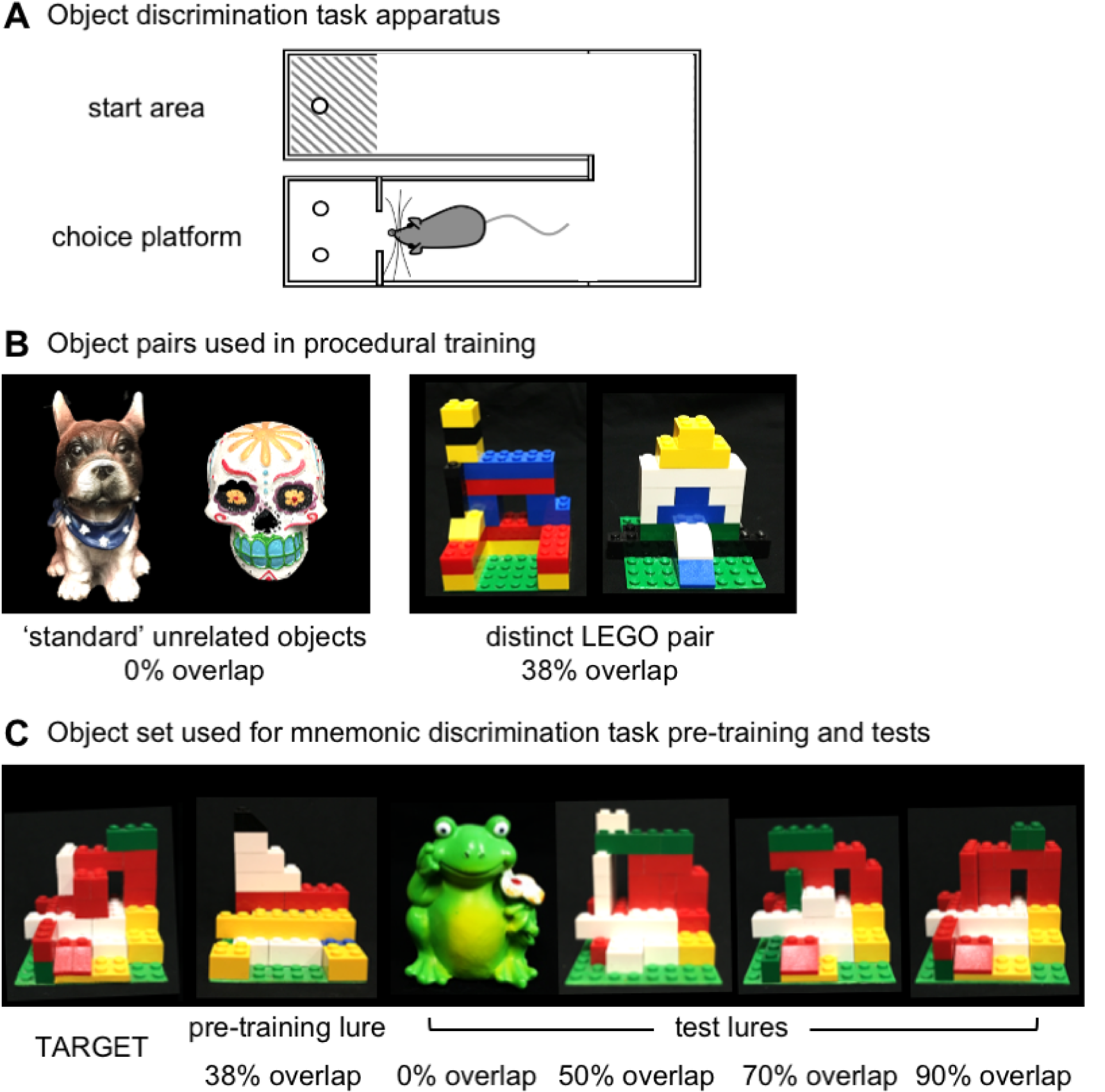
Rat mnemonic similarity task apparatus. **(A)** U-shaped maze used for the object-based mnemonic discrimination task. Rats were placed in the start area at the beginning of a session, then traversed to the choice platform where object stimuli were placed over food wells. After response selection, rats returned to the start area to initiate the next trial. Walls 25-30 cm in height surround the choice platform to prevent rats from visualizing objects as they are placed by the experimenter. **(B)** Object stimulus pairs used for procedural training. Rats were first trained with ‘standard’ unrelated junk objects that share 0% visible feature overlap, then with perceptually distinct LEGO objects that share 38% feature overlap. For each pair, one object served as the target (S+) and the alternate object as a lure (S-). The object serving as the target for each pair was counterbalanced across rats. (C) Object stimuli used for the rat mnemonic similarity task. For mnemonic discrimination tests, one object always served as the target (S+) for all rats (left; TARGET). Rats were pre-trained to identify this target with a single distinct LEGO lure that shared 38% feature overlap (pre-training lure). Rats were tested with four lure objects that ranged in similarity to the target: a distinct unrelated object with 0% feature overlap, and three LEGO lures that shared 50%, 70%, and 90% feature overlap. For a detailed description of feature overlap calculations see Johnson et al. 2017.

### Surgery

After procedural training, rats were implanted with bilateral guide cannulae targeting the prelimbic and infralimbic cortex. Isoflurane anesthesia was induced at 5%, then maintained at 1-3% (Isothesia, Henry Schein, Dublin, OH). A longitudinal incision was made to expose Bregma and Lambda and clean the skull surface. After confirming flat skull position in the stereotaxic apparatus, four stainless steel bone screws (#000-120, 1/8”; Antrin Miniature Specialties, Fallbrook, CA) were placed to anchor the head cap, two posterior to the coronal suture and two anterior to the lambdoid suture. Custom bilateral stainless steel guide cannulae (C252G DBL, 22G; P1 Technologies, Roanoke, VA) were positioned at AP +3.6 mm, ML ±0.9 mm, and DV -3.8 mm relative to Bregma at skull surface. Cannulae tips were positioned 1 mm dorsal to infusion sites, as microinjectors protruded 1 mm below this coordinate. Cannulae were secured to the skull and anchor screws with dental cement (Teets, Patterson Dental, Tampa, FL). Dust caps (303DC, P1 Technologies) protected cannulae from contamination. NSAIDS (Metacam, 1-2 mg/kg s.c., Boehringer Ingelheim, Vetmedica Inc., St. Joseph, MO) were administered for analgesia pre- and post-op. Rats were given a minimum recovery period of 1 week before resuming behavioral experiments.

### Mnemonic discrimination task

Following recovery, rats were pre-trained for mnemonic discrimination tests with a new pair of LEGO objects (**Fig.1C**). One object would serve as the target (S+) throughout testing and the other was a perceptually distinct lure (pre-training lure; S-) sharing 38% feature overlap, comparable to overlap of the distinct LEGO pair used in procedural training (**Fig.1B**). After reaching criterion of ≥81.3% correct responses on 2 consecutive training sessions, rats were given 2 days off before their first mnemonic discrimination test. Tests were carried out as previously described with the learned target and a set of lures that shared 0-90% feature overlap (**Fig.1C**) (Johnson et al., 2017; Burke et al., 2018b; Johnson et al., 2018). Each session consisted of 50 trials: 10 with an entirely distinct standard lure (frog figurine, 0% overlap), 10 with each of 3 perceptually similar LEGO lure objects, sharing 50, 70, and 90% feature overlap, and 10 with an identical copy of the target object. Trials with the copy of the target were included as a control condition, to verify rats were not using odor cues from the maze, objects, or food reward to guide response selection. Sessions were recorded with a camera mounted directly above the choice platform. Response latencies and response selection behavior were scored offline from recordings with custom software (Collector/Minion; Burke/Maurer Labs, Gainesville, FL).

### Intra-cerebral infusions

Tests proceeded every 3 days to avoid over-familiarization with lure objects and provide a wash-out period between drug infusions, as in prior studies (Bañuelos et al., 2014; McQuail et al., 2016; Johnson et al., 2018). Order of infusion conditions was pseudo-randomized across 3 test blocks in a Latin squares design, such that each rat received both a vehicle and a muscimol infusion within each test block. The GABA_A_ receptor agonist muscimol (1 mg/mL, Sigma-Aldrich, St. Louis, MO) or vehicle (0.9% sterile saline) were infused bilaterally to the prelimbic/infralimbic cortex (0.5 μL per hemisphere, 0.1 μL/min) 30 min prior to each test. Internal bilateral stainless steel microinjectors (C232I, 28G, P1 Technologies) protruding 1 mm below implanted guides were connected via polyethylene tubing (PE50, P1 Technologies) to 10 μL syringes (Hamilton, Franklin, MA) mounted in a microinfusion pump (Harvard Apparatus, Holliston, MA). Tubing was backfilled with sterile water. A 1-μL bubble was aspirated to create a barrier between backfill and infusate, and permit confirmation by visual inspection that the correct volume of infusate had been delivered. Microinjectors were left in place for 2 min after the infusion to allow dispersion of the drug. Rats were returned to the home cage for 30 min prior to beginning behavioral testing.

### Verification of cannulae placements and effects of muscimol

Positions of guide cannulae tips and effects of muscimol infusions on neuronal activity were verified at the conclusion of experiments, as in previous studies (Hernandez et al., 2017; Johnson et al., 2018). Briefly, effects of muscimol within the diffusion radius of the drug were assessed by visualizing cells expressing mRNA of the immediate-early gene *Arc* with fluorescence *in situ* hybridization (FISH). One week after completing infusions and mnemonic discrimination test blocks, rats received a bilateral intra-cerebral muscimol infusion (0.5 μL per hemisphere, 1 mg/mL) and were returned to their home cages for 30 min. Rats then performed a 10-min object discrimination epoch, were transferred to a glass bell jar for deep anesthesia with isoflurane, and were euthanized by rapid decapitation. In this behavioral epoch, the target object and lure object sharing 70% feature overlap (**Fig.1C**) were used as stimuli on all trials. Rats completed as many trials as possible within 10 min; this ranged from 16 to 28 trials (mean = 23, SD = 4.73). Brains were extracted and snap-frozen in chilled 2-methyl butane (Acros Organics, Fisher Scientific, Pittsburgh, PA). Tissue blocks were sectioned (20 μm) on a cryostat (Microm HM550, ThermoFisher Scientific, Waltham, MA), thaw-mounted on Superfrost Plus slides (Fisher Scientific) and stored at -80°C in sealed slide boxes. One rat prematurely lost their head cap and had to be euthanized; this rat was excluded from neural analyses of muscimol effects. Instead, this rat was perfused and cannulae placements were verified by imaging 40 μm sections with fluorescence microscopy (Keyence, Itasca, IL).

FISH for *Arc* mRNA was carried out as previously described (Hernandez et al., 2017, 2018b; Maurer et al., 2017; Johnson et al., 2018). Briefly, digoxigenin (DIG)-labeled riboprobes were generated with a commercial transcription kit (Riboprobe System, P1440, Promega, San Luis Obispo, CA) and DIG RNA labeling mix (Roche REF# 11277073910, MilliporeSigma, St. Louis, MO) from a 3.0 kb *Arc* cDNA template (provided by Dr. A. Vazdarjanova, Augusta University, Augusta, GA). Slides were hybridized and incubated with anti-digoxigenin-POD (Roche REF# 11207733910, MilliporeSigma). Labeled transcripts were visualized with Cy3 (TSA Cyanine 3 System, NEL744A001KT, PerkinElmer, Waltham, MA) and sections were counter-stained with DAPI. Low magnification stitched images were collected by fluorescence microscopy (Keyence) with a 2X objective. Cannulae placements were verified from these flyover images. Placements of implanted cannulae for all rats are shown in Figure 2A. In two slides adjacent to cannulae tracks, approximately 3.6-4.6 mm anterior to Bregma, high magnification z-stacks were captured with a 40X objective from prelimbic (PrL) and infralimbic (IL), anterior cingulate (Cg1), orbitofrontal (OFC), and motor (M1) cortices (**Fig.2B**). Proportions of neurons with *Arc* mRNA labeling in each region were determined by manually counting *Arc*-positive DAPI-labeled neuronal nuclei with ImageJ software and a custom plug-in, as previously described (Hernandez et al., 2017, 2018b; Maurer et al., 2017; Johnson et al., 2018).

**Figure 2.**
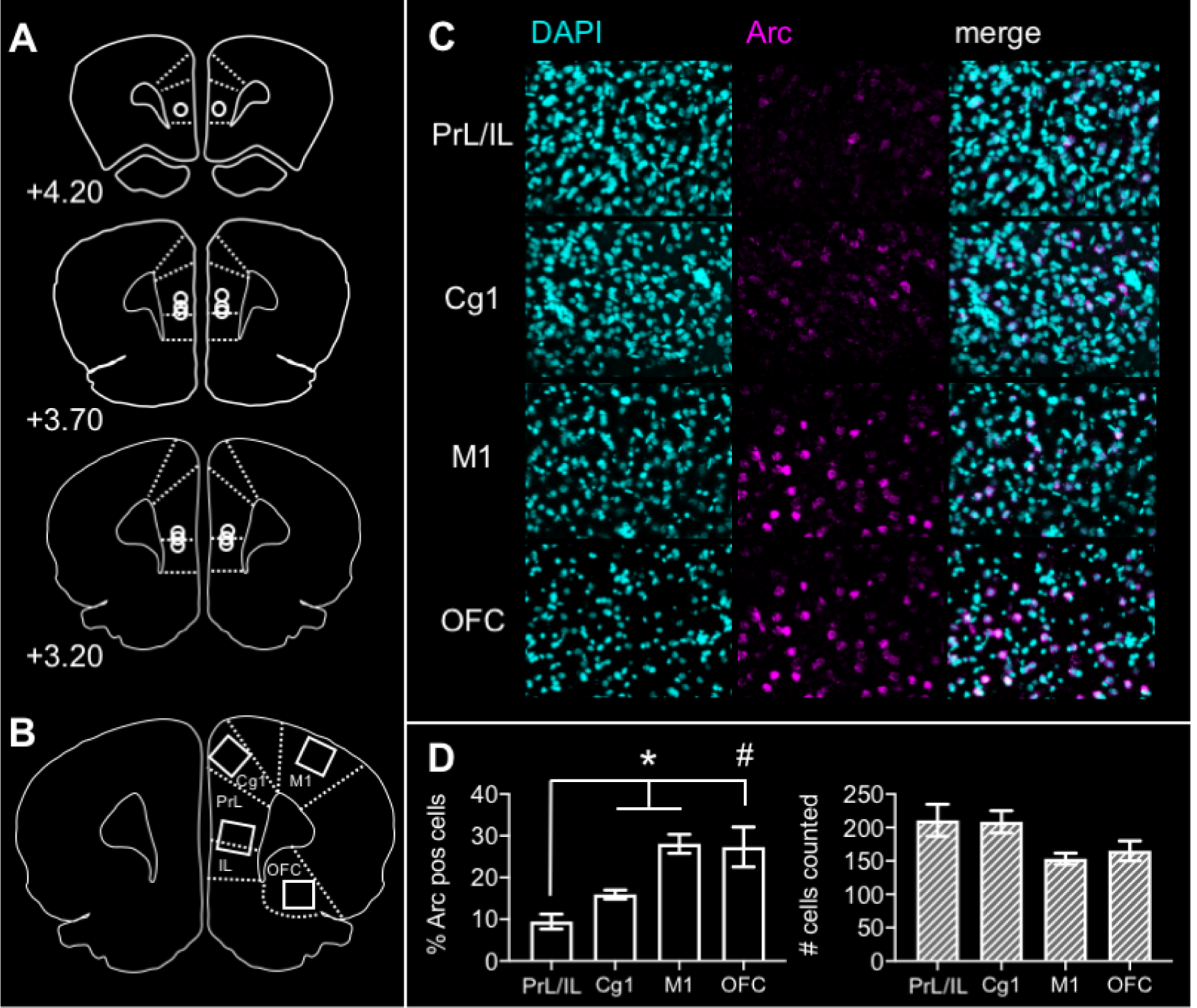
Verification of guide cannulae positions and neural effects of prefrontal cortical muscimol infusions. **(A)** Placements of microinjector tips are shown on representative prefrontal cortical sections for each rat from the study, AP +3.2-4.2mm relative to Bregma. **(B)** Regions of interest sampled from tissue sections adjacent to guide cannulae tracks. High magnification z-stacks were captured from prelimbic (PrL), infralimbic (IL), anterior cingulate (Cg1), orbitofrontal (OFC), and motor (M1) cortices. (**C**) Representative z-stacks from each region of interest labeled with the nuclear marker DAPI (cyan) and for mRNA of the neuronal activity-dependent immediate early gene *Arc* (magenta). **(D)** Proportion of neurons expressing *Arc* mRNA (left panel) and total number of neurons counted for the analysis (right panel) are shown for each region of interest. Following prefrontal muscimol infusions, the proportion of *Arc* mRNA-positive neurons in PrL/IL was significantly less than that present in Cg1 (contrasts; p = 0.016) and M1 (p = 0.003). The same trend was observed for PrL/IL relative to neurons in OFC (p = 0.059). Total number of cells counted did not differ across regions (main effect region; p = 0.095). * p<0.05,# p < 0.10.

### Statistical analyses

Analyses were performed with the Statistical Package for the Social Sciences (IBM SPSS) v25 for Mac OSX. Behavioral variables were analyzed with repeated measures ANOVAs, with test block, infusion condition, and lure object as within subjects factors, and within subjects or simple contrasts for post hoc analyses. Initial analyses were carried out with sex as a between subjects factor. No significant main effects of sex were detected for any behavioral measure, therefore sex was not included as an independent variable in subsequent analyses. When relevant, performance was compared to chance levels (i.e., 50% correct responses) with one-sample t-tests. Choice of statistical test was dictated by assumptions of normality, assessed with Shapiro-Wilk tests, and homogeneity of variances, assessed with Levene’s tests. Loss of several behavioral video recordings due to technical issues resulted in missing values for response latency data. These data were treated as independent samples by infusion condition and test block and were analyzed with non-parametric Mann Whitney U tests and Kruskal Wallis tests. P-values less than 0.05 were considered statistically significant, with Bonferroni corrections applied in cases where multiple comparisons were carried out.

## Results

### Verification of cannulae placements and neural effects of muscimol

Cannulae positions for all rats were located within the prelimbic and infralimbic (PrL/IL) cortex in the region spanning 3.20-4.20mm anterior to Bregma (**Fig.2A**). To confirm muscimol had the intended effect of inhibiting neural activity in PrL/IL, proportions of neurons expressing mRNA of the immediate-early gene *Arc* were determined from these regions (PrL/IL) and adjacent regions within the same coronal section, approximately 0.5 mm anterior to the cannulae placement (Cg1, M1, OFC; **Fig.2B**). Representative z-stacks from each region of interest are shown in Figure 2C. Percentages of *Arc*-positive neurons and total number of cells sampled in each region are shown in Figure 2D. Repeated measures ANOVA revealed a significant main effect of region (*F*(3,12) = 6.575, *p* = .007). Within subjects contrasts confirmed fewer *Arc*-positive neurons present in PrL/IL relative to adjacent cingulate cortex (Cg1; *F*(1,4) = 16.324, *p* = .016) and motor cortex (M1; *F*(1,4) = 44.473, *p* = .003). The difference in percent *Arc*-positive neurons in PrL/IL relative to orbitofrontal cortex was not statistically significant but followed the same trend (OFC; *F*(1,4) = 6.857, *p* = .059). The total number of cells counted did not differ across cortical regions (RM ANOVA main effect of region, *F*(3,12) = 2.668, *<u>p</u>* = .095).

### Procedural training

Figure 3 shows the number of incorrect trials completed by rats prior to reaching criterion in procedural training and pre-training with the target object used in the mnemonic discrimination task. Rats differed in amount of training required to reach criterion performance of >81% correct responses across object pairs (RM ANOVA main effect of pair, *F*(2,10) = 17.138, *p* = .001). Follow-up contrasts confirmed this effect was attributable to a greater number of trials completed in learning to distinguish the distinct LEGO object pair used in procedural training (distinct LEGO pair vs. standard objects, *p* = .01; vs. pre-training LEGO objects, *p* = .002). The fact that all rats learned to accurately discriminate the mnemonic discrimination target object and distinct pre-training lure within a comparable number of trials as the first standard object pair (contrast, *p* = .340) confirms surgical implantation of prefrontal cortical cannulae did not interfere with existing procedural knowledge of the discrimination task, nor impede rats’ abilities to learn the identity of a new target object.

**Figure 3.**
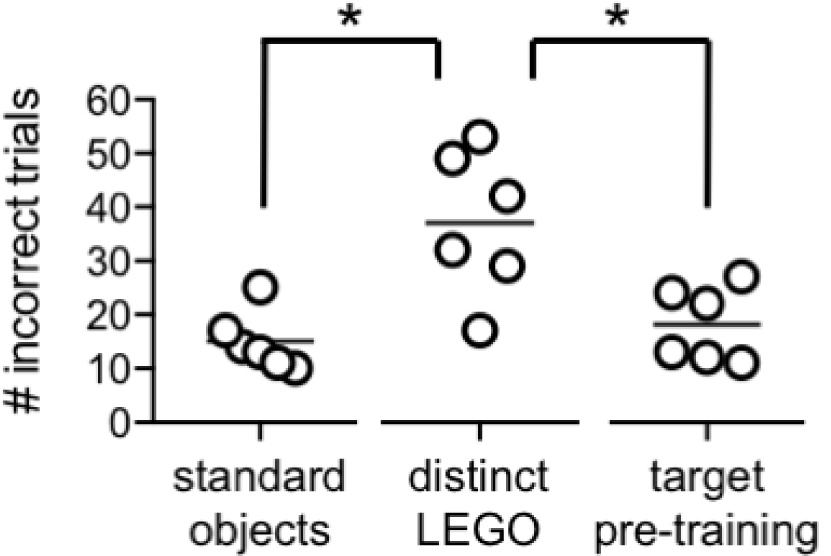
Procedural training data. Total number of incorrect trials completed prior to reaching a criterion performance level of ≥ 26/32 (≥ 81%) correct responses on each of 2 consecutive daily object discrimination training sessions. Data are shown for the standard unrelated object pair (0% feature overlap) and distinct LEGO pair (38% feature overlap) used in procedural training, and the LEGO target object and distinct lure (38% feature overlap) used after cannulae implantation for mnemonic similarity task pre-training. Total number of incorrect trials differed across object pairs (main effect object pair; p = 0.001). Rats required a greater number of trials to learn the distinct LEGO discrimination during procedural training than the standard object discrimination (contrasts; p = 0.01). However, rats then took fewer trials to learn the target and distinct LEGO lure in mnemonic discrimination pre-training after cannulae implantation (p = 0.002), confirming surgical implantation of guide cannulae did not interfere with rats’ abilities to learn a new lure or draw on procedural knowledge acquired pre-operatively.

### Prefrontal cortical muscimol impairs mnemonic discrimination performance

Muscimol or vehicle infusions were administered prior to mnemonic discrimination tests across three blocks in a Latin squares design (**Fig.4A**). Performance was first analyzed with infusion condition, test block, and lure object as within subjects factors. As in prior studies (Johnson et al., 2017, 2018), performance on control trials in which rats were presented with two identical copies of the target object (i.e., 100% overlap condition) did not differ across test blocks (RM ANOVA main effect of block; *F*(2,12) = 1.032, *p* = .386). Mean performance on these trials by block ranged from 47.1 – 54.3 (SD = 5.67 – 12.15) and did not differ from chance in blocks 1 and 2 (one-sample t-tests with hypothetical mean of 50% correct trials; block 1, *p* = .231, block 2, *p* = .766). With alpha levels corrected for multiple comparisons (.05 / 3 = .017), performance on these trials in block 3 also did not differ from chance (*p* = .045). Data from these trials were therefore excluded from subsequent analyses.

Percent correct responses on the mnemonic discrimination task collapsed across the three test blocks is shown in Figure 4B. Repeated measures ANOVA revealed that, overall, prefrontal cortical muscimol infusions impaired rats’ performance on the task (main effect of infusion condition; *F*(1,6) = 76.364, *p* < .0001). Consistent with previous studies (Johnson et al., 2018, 2017), performance also differed based on the similarity of lure objects to the target object (main effect of lure; *F*(3,18) = 34.720, *p* < .0001). A significant infusion condition x lure object interaction (*F*(3,18) = 4.294, *p* = .019) suggested that the degree to which muscimol impaired performance differed based on target-lure similarity. Follow-up within subjects contrasts showed relative to vehicle, muscimol impaired rats’ abilities to distinguish the target from similar lure objects (i.e. sharing 50% or more visible feature overlap; 50% overlap, *p* < .0001, 70% overlap, *p* < .0001, 90% overlap, *p* = .001). After correcting alpha levels for multiple comparisons (.05 / 4 = .013), the difference in performance on trials with the distinct standard object lure (0% feature overlap) was not statistically significant (*p* = .017).

**Figure 4.**
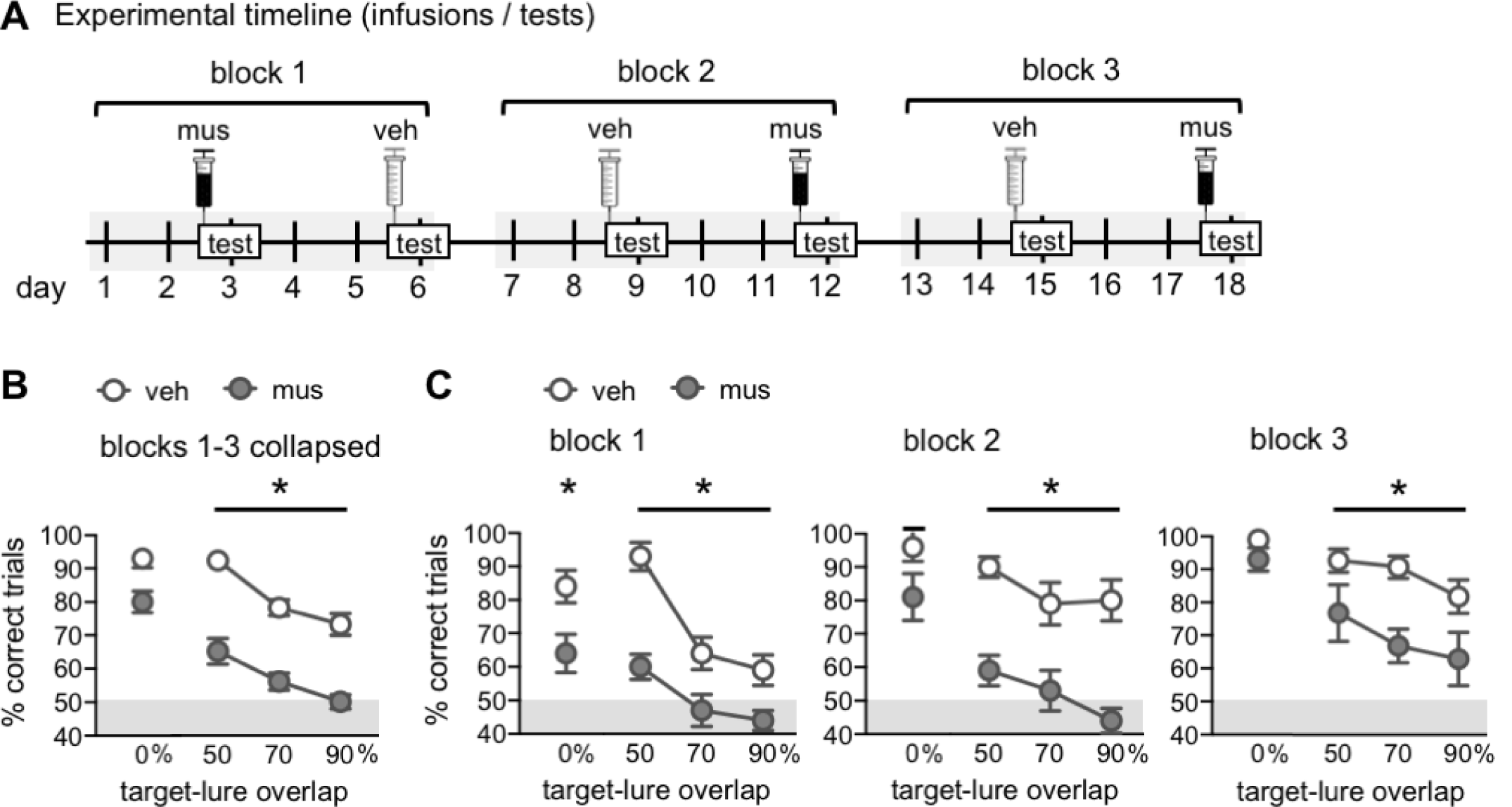
Effects of prefrontal cortical inactivation on mnemonic similarity task performance. **(A)** Experimental timeline. Muscimol (mus) or vehicle (veh) infusions were given prior to mnemonic discrimination tests across three blocks in a Latin squares design. Infusions/tests took place every 3 days to allow a sufficient period for drug wash-out. Order of infusion condition within each test block was counterbalanced across rats. **(B)** Mnemonic discrimination performance following vehicle (veh) and muscimol (mus) infusions, collapsed across test blocks 1-3. Performance is shown as mean percent (%) correct trials for trial types with each level of target-lure overlap (0%, 50%, 70%, and 90%). Prefrontal cortical muscimol infusions impaired mnemonic discrimination performance across test blocks (main effect infusion; p = < 0.001) but effects differed based on target-lure overlap (interaction infusion x lure; p = 0.019). Follow-up contrasts showed muscimol impaired discrimination on trials with LEGO lure objects with overlap of 50% (p < 0.001), 70% (p < 0.001), and 90% (p = 0.001). (**C**) Mnemonic discrimination performance following vehicle (veh) or muscimol (mus) infusions on each of test blocks 1-3. As in (B), performance is shown as mean % correct trials for each level of target-lure overlap. Prefrontal cortical muscimol infusions impaired discrimination in each test block (main effects infusion; block 1, p = 0.007, block 2, p = 0.001, block 3 = 0.012). Performance on trials with similar lures, sharing target-lure overlap of 50%, 70%, and 90% remained impaired across all blocks (simple effects; p’s > 0.106), whereas after vehicle infusions, performance on trials with lures sharing overlap of 70% and 90% significantly improved (p’s < 0.01). * p < 0.05.

Percent correct responses on tests of blocks 1-3 following vehicle versus muscimol infusions are shown in Figure 4C. In our previous studies, we have found the effects of hippocampal muscimol infusions on mnemonic discrimination performance can differ as rats gain experience with the task across test blocks (Johnson et al., 2018). A repeated measures ANOVA with test block, infusion condition, and lure object as within subject factors was performed to determine if this was also the case for prefrontal cortical muscimol infusions. This analysis revealed significant main effects of block (*F*(2,12) = 12.830, *p* = .001), infusion (*F*(1,6) = 76.364, *p* < .0001), and lure (*F*(3,18) = 34.720, *p* < .0001), a significant infusion x lure interaction (*F*(3,18) = 4.294, *p* = .019), and a significant 3-way interaction (*F*(6,36) = 2.565, *p* = .036). Interactions of block x infusion (*F*(2,12) = .348, *p* = .713) and block x lure (*F*(6,36) = 1.679, *p* = .155) were not statistically significant.

To clarify effects of task experience, separate repeated measures ANOVAs were run on data from each test block, with lure object and infusion condition entered as within subjects factors. As in prior studies (Burke et al. 2018; Johnson et al., 2018, 2017), significant main effects of lure were observed across all test blocks (*F*’s > 11.663, *p*’s < .0001) confirming that performance across 3 blocks of discrimination tests remains contingent on target-lure overlap. Significant main effects of prefrontal infusion condition were detected in block 1 (*F*(1,6) = 16.241, *p* = .007), block 2 (*F*(1,6) = 43.070, *p* = .001), and block 3 (*F*(1,6) = 12.590, *p* = .012). Significant lure x infusion condition interactions were observed in block 1 (*F*(3,18) = 4.100, *p* = .022) and block 3 (*F*(3,18) = 5.210, *p* = .009), but not block 2 (*F*(3,18) = 1.957, *p* = .157). Additional repeated measures ANOVAs with lure object and test block as within subjects factors, holding infusion condition constant, showed significant main effects of lure (*F*’s > 22.041, *p*’s < .0001) and of block (vehicle, *F*(2,12) = 8.605, *p* = .005; muscimol, *F*(2,12) = 4.702, *p* = .031). However, a significant lure x block interaction was only observed on tests following vehicle infusions (*F*(6,36) = 3.022, *p* = .017), not following muscimol infusions (*F*(6,36) = 1.251, *p* = .304).

Simple effects analyses showed performance on trials with the distinct object lure (0% overlap) improved across blocks, for vehicle (*F*(2,12) = 6.326, *p* = .013) and for muscimol infusion conditions (*F*(2,12) = 6.423, *p* = .013). Comparably, after vehicle infusions, performance on trials with similar lures improved across test blocks. This was the case for trials with 70% overlap (*F*(2,12) = 6.923, *p* = .01) and 90% overlap (*F*(2,12) = 6.235, *p* = .014), but not 50% overlap (*F*(2,12) = .673, *p* = .528), because rats’ performance remained high on these trials across blocks. After muscimol infusions, performance on trials with similar lures did not improve across blocks. Simple effects did not show statistically significant differences for trials with 50% overlap (*F*(2,12) = 1.864, *p* = .197), 70% overlap (*F*(2,12) = 1.511, *p* = .260), or 90% overlap (*F*(2,12) = 2.717, *p* = .106).

### Prefrontal cortical muscimol causes rats to default to a side-biased strategy

Previous studies (Hernandez et al., 2015, 2017; Johnson et al., 2017; Burke et al., 2018b; Johnson et al., 2018) have found that rats default to selecting an object on a particular side (regardless of features) when learning forced-choice discrimination tasks. This side-biased strategy is particularly prominent in aged rats (Hernandez et al. 2015; Johnson et al. 2017) and young adult rats in which connectivity across perirhinal cortex, hippocampus, and/or the prefrontal cortex is experimentally disrupted (Lee and Solivan, 2008; Jo and Lee, 2010a, 2010b; Hernandez et al., 2017; Burke et al., 2018b; Johnson et al., 2018). To determine if this was the case in the current study, side bias indices were calculated as the absolute value of (total number of left well responses – total number of right well responses) / total number trials. Figure 5A shows these values collapsed across trial types for each of the tests given in blocks 1-3. As in prior studies, muscimol caused rats to have a greater side bias index, relative to vehicle (RM ANOVA, main effect of infusion condition, F(1,6) = 24.449, p = .003). The side bias index did not vary across tests (main effect of block, F(2,12) = 2.190, p = .155; infusion x block interaction, F(2,12) = .013, p = .702). However, within subjects contrasts showed that following vehicle infusions, side bias decreased from block 1 to block 3 (F(1,6) = 14.583, p = .009), while after muscimol infusions, side bias remained the same from block 1 to block 3 (F(1,6) = .373, p = .564).

**Figure 5.**
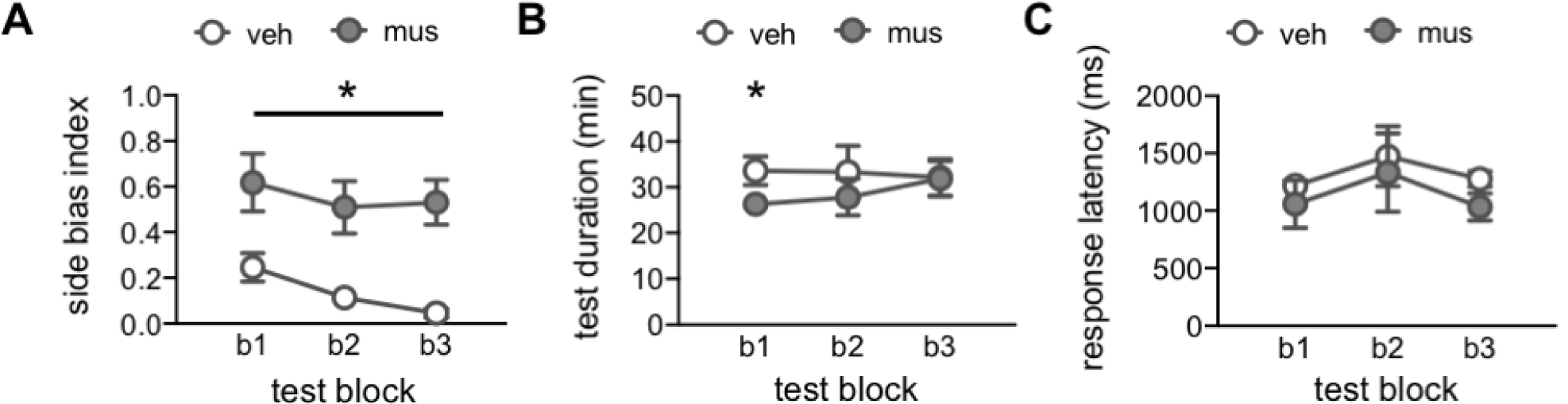
Side bias index, test session duration, and response selection latency. **(A)** Side bias index values computed for each test block (blocks 1-3; b1, b2, b3) following vehicle (veh) or muscimol (mus) infusions. Side bias index is calculated as the absolute value of (total number of left well responses -total number of right well responses) *I* total number trials, such that a value of 0 reflects no side bias and an equal number of responses made to left and right wells, and a value of 1 reflects a complete bias to either the left or right side well. Muscimol infusions led rats to have a greater side bias across all test blocks (main effect infusion; p = 0.003). **(B)** Session durations (minutes; min) by infusion condition across test blocks. Prefrontal muscimol infusions did not slow behavioral responding on the mnemonic discrimination task, but rather led rats to complete test sessions in a shorter period of time relative to vehicle infusions (main effect infusion; p = 0.035). (C) Response selection latencies (milliseconds; ms) for correct trials by infusion condition across test blocks. Prefrontal muscimol infusions did not alter response latencies relative to vehicle infusions on any test block (p’s > 0.085). * p < 0.05.

### Prefrontal cortical muscimol does not slow behavior or response selection

An important consideration in interpreting the effects of temporary neuronal inactivation is the potential for muscimol to disrupt motor function or motivation, impeding task performance. Analyses confirmed that all rats completed full test sessions of 50 trials after receiving prefrontal cortical infusions of muscimol. The duration of test sessions following vehicle vs. muscimol infusions is shown in Figure 5B. A repeated measures ANOVA with test block and infusion condition as within subjects factors revealed a significant main effect of infusion (F(1,4) = 9.898, p = .035), but no main effect of block or block x infusion interaction (F’s < .624, p’s > .560). Within subjects contrasts showed the duration of test sessions differed between infusion conditions only on block 1 (p = .017, blocks 2-3, p’s > .173). Critically, rats completed their test sessions *faster* following prefrontal muscimol infusions on this first test block, relative to vehicle infusions.

To determine if this speeding of test completion was attributable to faster response selection, response latencies were scored on video recordings of each test session. As in prior studies (Johnson et al., 2017; Burke et al., 2018b; Johnson et al., 2018), response latencies are defined as the time elapsed between the point at which the rat’s nose crossed the threshold of the choice platform and the point at which the rat initiated displacement of one of the objects on the choice platform. Response latencies for correct responses to rats’ inherent biased side are shown in Figure 5C. Latencies for incorrect responses and responses to rats’ inherent non-biased side are excluded because of missing data on tests following muscimol infusions; in most cases, rats made no responses to their non-biased side after prefrontal muscimol infusions. Because some video recordings were lost due to technical issues, response latency data from each block and drug condition were treated as independent samples and analyzed with non-parametric tests. The effect of infusion condition on correct response latencies for each test block was assessed with Mann Whitney U tests (asymptotic significance, 2-tailed). Latencies did not differ based on infusion condition in block 1 (mean latency vehicle = 1218 ms versus muscimol = 1058 ms; U = 11, n_1/2_ = 7, p = .085), block 2 (mean latency vehicle = 1475 ms versus muscimol = 1332 ms; U = 13, n_1/2_ = 6, p = .423), or block 3 (mean latency vehicle =1276 ms versus muscimol = 1033 ms; U = 4.0, n_1_ = 4, n_2_ = 5, p = .142). The effect of test block on correct response latencies was also assessed with Kruskal-Wallis tests. Latencies did not differ by block after vehicle infusions (H(2) = 0.520, p = 0.771) or after muscimol infusions (H(2) = 0.500, p = 0.779).

## Discussion

In the current study, we document a novel role for prefrontal cortical activity in the mnemonic discrimination of similar object stimuli. Our main finding is that prefrontal cortical inactivation impairs rats’ abilities to distinguish a previously learned target object (S+) from lure objects (S-) that share overlapping visible features. The mnemonic discrimination of similar stimuli has long been associated with hippocampal function, in part based on models suggesting pattern separation computations arise from hippocampal circuits (Marr, 1971; McNaughton and Morris, 1987; Treves and Rolls, 1992; O’Reilly and McClelland, 1994; Yassa and Stark, 2011; Santoro, 2013; Kesner and Rolls, 2015; Knierim and Neunuebel, 2016; Leal and Yassa, 2018), and based on functional neuroimaging data from humans showing activation of hippocampal dentate gyrus (DG)/CA3 is associated with mnemonic discrimination performance across adulthood (Kirwan and Stark, 2007; Bakker et al., 2008; Lacy et al., 2011; Yassa and Stark, 2011; Motley and Kirwan, 2012; Reagh and Yassa, 2014; Doxey and Kirwan, 2015; Reagh et al., 2018). In animal models, hippocampal lesions impair discrimination of similar spatial locations (Gilbert et al., 1998, 2001; Morris et al., 2012; Oomen et al., 2015), yet until recently, the requirement of intact DG and CA3 activity for discrimination of similar visual stimuli – akin to those presented in human mnemonic similarity tasks – had not been investigated. Using an object-based rat mnemonic similarity task (Johnson et al., 2017), we therefore sought to test whether reversibly disrupting neural activity in the DG and CA3 selectively interfered with discrimination of a known target from similar lures (Johnson et al., 2018). Results of this set of experiments initially showed, when collapsing performance across three repeated tests, muscimol infusions in the dorsal DG and CA3 had no effect on mnemonic discrimination of a known target from distinct or similar lures (Johnson et al., 2018). This observation is distinct from the results of the current experiments that showed, when collapsing across three repeated tests, muscimol infusions in the prefrontal cortex impaired discrimination of the known target from all distinct and similar lures (**Fig.4B**). To clarify the effect of hippocampal muscimol infusions, additional experiments revealed that disruption of DG and CA3 activity impaired mnemonic discrimination only on a first test when lure objects were novel (Johnson et al., 2018). On subsequent tests, when the lure stimuli were familiar, hippocampal muscimol infusions were found to not significantly impair rats’ performance. Analysis of behavioral performance across tests showed muscimol infusions in the prefrontal cortex consistently impaired discrimination of the known target from lures sharing 50-90% feature overlap (**Fig.4C**). Further, performance on trials with these lures did not improve across tests with muscimol infusions, despite performance improving on trials with 70-90% overlap across tests with vehicle infusions (**Fig.4C**). Together these data indicate that both the hippocampus and the prefrontal cortex are critical for mnemonic discrimination performance but the time course of involvement and modulation by novelty of lures is dissociable.

### Prefrontal cortical contributions to hippocampal-dependent memory

There are several potential interpretations of the dissociable effects of dorsal hippocampal versus prefrontal cortical muscimol infusions. A first possibility is that, as task stimuli and procedures become increasingly familiar, cortical regions can compensate for hippocampal dysfunction. Neural activity across prefrontal cortical regions could therefore account for rats’ abilities to overcome DG and CA3 dysfunction in the mnemonic discrimination task once they have gained experience with lure stimuli (Johnson et al., 2018). There are several bases for this idea. Well-known models of systems-level memory consolidation argue hippocampal activity is most required for learning and formation of new memory representations, and that these representations become distributed across prefrontal cortical regions over time (Frankland and Bontempi, 2005; Eichenbaum, 2017; Takehara-Nishiuchi, 2020). On the other hand, it is increasingly acknowledged that specific functions or phases of memory processing cannot necessarily be ascribed to specific brain circuits, and that compensation for damage to a given region is possible due to dynamic and distributed processing (Fanselow, 2010; Ash and Rapp, 2014; Rubin et al., 2017; Burke et al., 2018a). Our current results suggest the prefrontal cortex is a critical node in a distributed cortical-hippocampal circuit engaged by mnemonic discrimination.

Another study using contextual fear conditioning in rats points to the prefrontal cortex as a site of compensation for hippocampal dysfunction (Zelikowsky et al., 2013). In these experiments, dorsal hippocampal lesions led to retrograde – but not anterograde – impairments in expression and renewal of contextually conditioned fear memory (Zelikowsky et al., 2013). However, dual lesions of dorsal hippocampus and the infralimbic or prelimbic cortex, or lesions of only the infralimbic cortex, impaired new contextual fear learning, discrimination, and renewal (Zelikowsky et al., 2013). Further, rats with lesions to only the dorsal hippocampus showed increased neural activation in the prelimbic cortex associated with unimpaired contextual fear memory retrieval (Zelikowsky et al., 2013). What is most intriguing about these findings with respect to dynamic processing is that the medial prefrontal cortical regions compensating for hippocampal dysfunction do not share direct projections with the dorsal hippocampal sub-regions that were lesioned. The prelimbic and infralimbic cortices receive monosynaptic projections from ventral hippocampus, but mainly interface with the hippocampus through multi-synaptic reciprocal connections via entorhinal and perirhinal cortices, as well as the thalamic nucleus reuniens (Swanson, 1981; Swanson and Köhler, 1986; Burwell and Amaral, 1998; Vertes, 2004; Cenquizca and Swanson, 2007; Hoover and Vertes, 2007; Agster and Burwell, 2009; Hwang et al., 2018). This emphasizes the distributed nature of circuit activity that occurs during multiple forms of memory processing traditionally linked to hippocampal function.

The current data indicate that a larger distributed network is critical for mnemonic similarity task performance, which is consistent with human imaging data. In young adults, mnemonic discrimination abilities correlate with activation of lateral entorhinal and perirhinal cortices to the same extent as activation of hippocampal CA3/DG (Reagh and Yassa, 2014). In contrast, in older adults, mnemonic discrimination impairments are associated with decreased activity in lateral entorhinal and perirhinal cortices, increased activity in CA3/DG (Bakker et al., 2012, 2015; Ryan et al., 2012; Berron et al., 2018, 2019; Reagh et al., 2018), and loss of perforant path white matter integrity (Yassa et al., 2010; Bennett et al., 2015; Bennett and Stark, 2016). These human data suggest entorhinal and perirhinal regions provide a hub through which neural activity coordinates mnemonic discrimination functions; however, few studies have examined activation in non-temporal cortical regions during these tasks. In animal models, several studies have directly tested this possibility using other hippocampal-dependent tasks. Disconnection of the perirhinal cortex from medial prefrontal cortex by infusing muscimol in contralateral hemispheres impaired recall of previously learned object-place paired associations (Hernandez et al., 2017) that depend on the hippocampus (Lee and Solivan, 2008; Jo and Lee, 2010a). In fact, blockade of prefrontal-perirhinal communication impaired object-place memory to the same extent as inactivation of medial prefrontal cortex or perirhinal cortex on their own (Hernandez et al., 2017). Similarly, reversible inactivation of neurons in the perirhinal cortex or thalamic nucleus reuniens that receive medial prefrontal cortex input, with AAV-mediated expression of the inhibitory DREADD hM4Di, impaired memory for previously learned odor sequences (Jayachandran et al., 2019). In this case, deficits produced by selectively targeting neurons that received prefrontal input were more profound that those resulting from direct inactivation of the medial prefrontal cortex itself (Jayachandran et al., 2019). The current study is the first, to our knowledge, to directly test involvement of prefrontal cortical activity in the mnemonic discrimination of similar object stimuli, and thus we have not yet explored how prefrontal communication with parahippocampal cortices influences rats’ performance on the task. The recent observation that unilateral transection of perforant path input from the lateral entorhinal cortex to the dorsal DG and CA3 selectively impairs young adult rats’ abilities to distinguish a known target object from similar lures (Burke et al., 2018b), suggests that lateral entorhinal cortical activity is also necessary for rats’ performance on the task. Additionally, it is well established that lesion or inactivation of the perirhinal cortex impairs object recognition and discrimination in rats and monkeys (Murray and Richmond, 2001; Bussey et al., 2002, 2003; Winters and Bussey, 2005; Bartko et al., 2007a, 2007b; Winters et al., 2010) and perirhinal neural activity can reflect perceptual differences in object stimuli that share feature overlap, as well as their mnemonic significance (Ahn and Lee, 2017). Thus, it logically follows that communication across parahippocampal and prefrontal cortices supports mnemonic discrimination of similar stimuli. This area remains open for future investigation.

### Prefrontal cortical contributions to high-demand tasks

A second potential interpretation of the current finding that prefrontal cortical activity is needed for mnemonic discrimination, irrespective of experience with similar lure objects, is that frontal regions are recruited during tasks that impose a high cognitive load. Mnemonic discrimination requires working memory (Hales et al., 2015), retention of stimulus representations across delays, resolution of interference across similar stimuli, and flexibility in selecting an appropriate response strategy. Thus, it is likely frontal cortical function plays a significant role in mnemonic discrimination, yet few studies have examined frontal cortical activity as it relates to discrimination abilities.

Until recently, only one human neuroimaging study had specifically addressed whether prefrontal and other neocortical regions show activation consistent with pattern separation and/or completion functions (Pidgeon and Morcom, 2016). During initial presentation of image stimuli, activation in response to parametrically varied features of image stimuli was consistent with pattern separation in occipitotemporal, lateral prefrontal, and right parietal cortical regions (Pidgeon and Morcom, 2016). Additionally, activation of occipital and lateral prefrontal regions during initial encoding was associated with accurate mnemonic discrimination in the test phase 24 h later, suggesting these cortical regions assist in generating distinct representations of similar stimuli to serve later memory (Pidgeon and Morcom, 2016). A subsequent human study tested the effect of unilaterally disrupting neural activity in ventromedial prefrontal cortex with repetitive transcranial magnetic stimulation (rTMS) on mnemonic discrimination performance (Wais et al., 2018). These experiments showed rTMS delivered to the prefrontal cortex, but not angular gyrus, impaired discrimination performance in young adult participants (Wais et al., 2018). In contrast, two recent functional neuroimaging studies examining hippocampal and neocortical recruitment during mnemonic similarity task performance found, as previously, activation patterns in hippocampal CA3/DG and perirhinal cortex were associated with accurate mnemonic discrimination (Klippenstein et al., 2020; Stevenson et al., 2020). However, while a similar pattern was observed in occipital cortex, this activity did not predict mnemonic discrimination performance to the same extent as CA3/DG (Klippenstein et al., 2020). Disparities among these human imaging findings may relate to the timing of stimulus presentations or to differences in fMRI analyses.

Prefrontal cortical involvement in mnemonic discrimination abilities has also not been widely investigated in aging. In one study, the same analyses that revealed decreased perforant path fiber integrity correlates with age-related mnemonic discrimination impairments also found loss of hippocampal cingulum white matter integrity to have the same consequence (Bennett and Stark, 2016). Given the hippocampal cingulum provides widespread connections from medial temporal lobe to frontal regions (Bubb et al., 2018), this could suggest individuals with deterioration of these white matter tracts are more prone to impairments because of lack of connectivity to support neural compensation. A more recent study found a similar loss of integrity of the parahippocampal cingulum in older adults with mild cognitive impairment, which correlated with decreased entorhinal cortical thickness and functional connectivity between hippocampal and posterior frontal cortical regions (Berron et al., 2020). While mnemonic discrimination abilities were not assessed in this study population, reduced hippocampal-cortical connectivity also associated with elevated cerebrospinal fluid levels of phosphorylated tau and amyloid-β (Berron et al., 2020). Given mnemonic discrimination impairments are observed in older adults with early signs of dementia or Alzheimer’s pathology (Bakker et al., 2012, 2015; Stark et al., 2013; Reagh et al., 2014; Berron et al., 2019), it is likely frontal-cortical disconnection in aging and its exacerbation by neurodegenerative pathology contribute to reduced abilities to perform well on high-demand tasks such as the distinction of similar stimuli from memory.

The role of prefrontal cortical regions in supporting performance on tasks with high cognitive demand has been more directly investigated in animal models. While effects of disrupting prefrontal cortical activity on mnemonic discrimination of objects had not previously been tested, infusion of the NMDA antagonist AP5 in the medial prefrontal cortex impaired mnemonic discrimination of both distinct and similar spatial locations in a touchscreen-based trial-unique delayed nonmatch-to-location (TUNL) task in rats (Davies et al., 2017). In contrast, ibotenic acid lesions of the medial prefrontal cortex did not impair spatial discriminations on the same touchscreen TUNL task in a second study in rats (McAllister et al., 2013). However, prefrontal lesions did impair performance when a post-sample delay or high interference conditions were introduced (McAllister et al., 2013), consistent with a role for prefrontal cortex in supporting performance when greater task demands are present. Additional data from hippocampal-dependent tasks in rats support the same conclusion. Lesions or reversible inactivation of the prelimbic cortex impair spatial working memory across long, but not brief, delays (Dunnett, 1990; Seamans et al., 1995; Floresco et al., 1997; Churchwell and Kesner, 2011), and muscimol infusions in the medial prefrontal cortex impair odor memory or match-to-sample performance under conditions with high interference across stimuli (Peters et al., 2013; Peters and Smith, 2020). Collectively, results of these experiments support an interpretation that prefrontal cortical activity is necessary to support performance on hippocampal-dependent tasks that pose a high cognitive demand, whether by requiring distinction of similar stimuli, imposing a working memory load, or increasing interference across trials.

## Conclusions

The results of this study demonstrate that disrupting neural activity in the prefrontal cortex impairs mnemonic discrimination of a known target object from similar lures in young adult rats, across repeated tests and irrespective of prior experience with task stimuli. This finding is in contrast to prior documented effects of disrupting neural activity in the hippocampal dentate gyrus and CA3, which impaired mnemonic discrimination on a first test when lure stimuli were novel, but not on subsequent tests as lure stimuli became increasingly familiar (Johnson et al., 2018). These findings highlight contributions of prefrontal cortical activity and, potentially, prefrontal-parahippocampal cortical connectivity to supporting memory processes considered to be primarily hippocampal-dependent. Furthermore, given sensitivity of mnemonic discrimination tasks to age-related decline, our results emphasize the importance of considering how hippocampal dysfunction linked to discrimination impairments influences prefrontal and parahippocampal cortical circuit activity, potential for compensation across prefrontal and parahippocampal cortical networks, and broader cognitive outcomes.

## Data Availability Statement

The data that support the findings of this study are available from the corresponding author upon reasonable request.

## Acknowledgements

We thank Edward ‘Sam’ Eusanio and Meena Ravi for help with behavioral video scoring. We also thank Abbi Hernandez and Katelyn Lubke for assistance with collecting and processing tissue for histology. This work was carried out with funds from the Evelyn F. McKnight Brain Research Foundation (SAJ, SNB), the National Institute on Aging K99 AG058786 (SAJ), R01 AG049722 (SNB), and a McKnight Brain Institute Fellowship (SAJ).

